# Hierarchical heuristic species delimitation under the multispecies coalescent model with migration

**DOI:** 10.1101/2023.09.10.557025

**Authors:** Daniel Kornai, Tomáš Flouri, Ziheng Yang

**Affiliations:** Department of Genetics, Evolution and Environment, University College London, UK

**Keywords:** bpp, genealogical divergence index, gene flow, multispecies coalescent, species delimitation, giraffes, milksnakes, sunfish

## Abstract

The multispecies coalescent (MSC) model accommodates genealogical fluctuations across the genome and provides a natural framework for comparative analysis of genomic sequence data to infer the history of species divergence and gene flow. Given a set of populations, hypotheses of species delimitation (and species phylogeny) may be formulated as instances of MSC models (e.g., MSC for one species versus MSC for two species) and compared using Bayesian model selection. This approach, implemented in the program bpp, has been found to be prone to over-splitting. Alternatively heuristic criteria based on population parameters under the MSC model (such as population/species divergence times, population sizes, and migration rates) estimated from genomic sequence data may be used to delimit species. Here we extend the approach of species delimitation using the genealogical divergence index (*gdi*) to develop hierarchical merge and split algorithms for heuristic species delimitation, and implement them in a python pipeline called hhsd. Applied to data simulated under a model of isolation by distance, the approach was able to recover the correct species delimitation, whereas model comparison by bpp failed. Analyses of empirical datasets suggest that the procedure may be less prone to over-splitting. We discuss possible strategies for accommodating paraphyletic species in the procedure, as well as the challenges of species delimitation based on heuristic criteria.

## Introduction

Delineation of species boundaries is important for characterizing patterns of biological diversity, especially during the current global changes in climate and environment. Traditionally, species were identified and distinguished using morphological characteristics. Genetic data are informative about many processes related to species delimitation and identification, including population identities, interspecific hybridization and gene flow, and phylogenetic relationships and divergence times among the populations (Bateson, 1909) (see, e.g., Fujita *et al*., 2012 for review). Early methods that use genetic data to identify and delimit species relied on simple genetic-distance cutoffs (such as the ‘3x’, ‘4x’, or ‘10x’ rules), requiring interspecific divergence to be a few times greater than intraspecific diversity (Hebert *et al*., 2003, 2004), or reciprocal monophyly in gene trees (Baum and Shaw, 1995) (see, e.g., Sites and Marshall, 2003 for a review). However, such criteria are too simplistic as they do not accommodate polymorphism in the ancestral populations and incomplete lineage sorting (Hudson and Turelli, 2003) or uncertainties in gene-tree reconstruction (Knowles and Carstens, 2007; Yang and Rannala, 2017).

The multispecies coalescent (MSC) model (Rannala and Yang, 2003) provides a framework for analysis of genomic sequence data from closely related species or populations to infer the order and timings of species/population divergences. Likelihood-based implementations of the MSC accommodate incomplete lineage sorting and stochastic variation in gene trees (so that reciprocal monophyly is not needed) as well as phylogenetic uncertainties at each locus (so that one does not have to rely on inferred gene trees), making it possible to infer population histories even when there is widespread incomplete lineage sorting and very little phylogenetic information at every locus (Xu and Yang, 2016; Jiao *et al*., 2021). The MSC model has also been extended to accommodate gene flow between species or populations, assuming either introgression (pulse of gene flow) or migration (continuous gene flow over an extended time period) (Jiao *et al*., 2021; Hibbins and Hahn, 2022), leading to MSC-with-introgression (MSC-I) and MSC-with-migration (MSC-M) models (Flouri *et al*., 2020, 2023). As interspecific hybridization/introgression appears to occur commonly in both plants and animals (e.g., *Arabidopsis*, Arnold *et al*., 2016; *Anopheles* mosquitoes, Fontaine *et al*., 2015; *Panthera* cats, Figueiro *et al*., 2017; and Hominins, Nielsen *et al*., 2017), it may be important to consider explicitly gene flow in species delimitation.

Given a set of populations, different species delimitations correspond to different ways of merging populations into the same species. Each species delimitation, together with the phylogeny for the delimited species, can be formulated as an instance of the MSC model and fitted to genomic sequence data sampled from the extant species or populations. Competing models of delimitation can then be compared via Bayesian model selection (i.e., using ior model probabilities or Bayes factors) (Yang and Rannala, 2010). In the Bayesian program bpp, this is accomplished by using a Markov chain Monte Carlo (MCMC) algorithm to calculate the posterior probabilities different MSC models (Yang and Rannala, 2010, 2014; Yang, 2015; Flouri *et al*., 2018). In simulations (Luo *et al*., 2018), bpp showed lower rates of species overestimation and underestimation than the generalized mixed Yule-coalescent method (Pons *et al*., 2006; Fujisawa and Barraclough, 2013) or the Poisson tree process method (Zhang *et al*., 2013). In empirical datasets, bpp was effective in identifying cryptic species among ancient lineages. For example, Ramirez-Reyes *et al*. (2020) identified 13 new species of leaf-toed geckoes in a lineage that diverged 30 Ma.

While being effective in detecting cryptic sympatric species, the approach of model selection implemented in bpp has often been noted to over-split, identifying more lineages as distinct species than many other methods, especially when applied to geographical populations or races (Sukumaran and Knowles, 2017). For example, Campillo *et al*. (2020) analyzed 99 population pairs in the genus *Drosophila* and found that bpp identified 80 pairs as distinct species, whereas reproductive isolation was identified in only 69 pairs. Similarly, Bamberger *et al*. (2022) examined 48 *Albinaria cretensis* land snail populations, and found that morphological classifications suggested 3–9 species while bpp suggested 45–48. Barley *et al*. (2018) simulated multiple populations from a single species that exhibits population structure and isolation by distance, and found that bpp delimits geographically separated populations as distinct species. Those results suggest that the lineages identified by bpp sometimes correspond to populations rather than species (Chambers and Hillis, 2020). A number of authors have expressed concerns about the apparent over-splitting of bpp (MacGuigan *et al*., 2021).

Rather than treating species delimitation as a model-selection problem, an alternative approach is to define species status using an empirical criterion based on parameters that characterize the history of population divergence and gene flow, such as the population split time (*τ*), (effective) population sizes (*θ* _*A*_, *θ*_*B*_), and (effective) migration rates (*M*_*AB*_, *M*_*BA*_). Those parameters can be estimated under the MSC from genomic data, with the stochasticity of the coalescent process and the phylogenetic uncertainty in genealogical trees accommodated. Jackson *et al*. (2017) introduced a criterion called the *genealogical divergence index* (*gdi*), by considering the probability that two sequences sampled from *A* (*a*_1_ and *a*_2_) coalesce before either coalesces with a sequence (*b*) sampled from *B* (fig. 1). When *a*_1_ and *a*_2_ coalesce first, the resulting gene tree has the topology *G*_1_ = ((*a*_1_, *a*_2_), *b*). Let its probability be *P*_1_ = ℙ (*G*_1_). In the case of no gene flow between *A* and *B*, this is given as

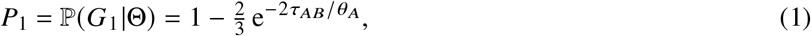

where the parameter vector is Θ = (*τ*_*AB*_, *θ* _*A*_, *θ*_*B*_, *θ*_*R*_), with *τ*_*AB*_ = *T*_*AB*_*μ* and *θ* _*A*_ = 4*N*_*A*_*μ*, where *T*_*AB*_ is the population split time in generations, *N*_*A*_ is the population size of *A*, and *μ* is the mutation rate per site per generation. Thus *τ* and *θ* _*A*_ measure the population split time and population size, respectively, both in expected numbers of mutations per site.

**Figure 1:**
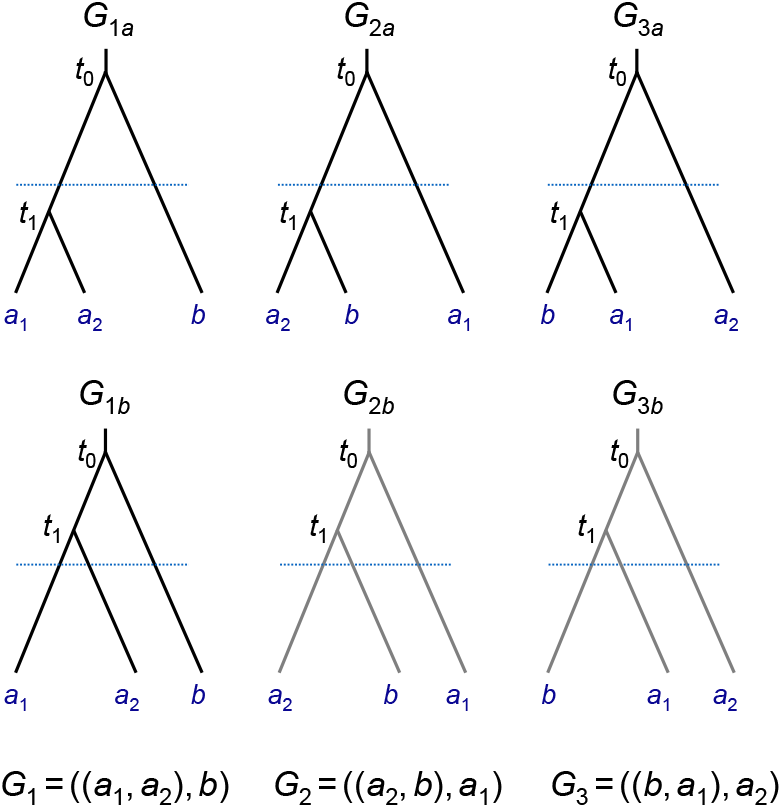
Three possible gene trees for a locus with two *A* sequences and one *B* sequence: *G*_1_ = ((*a*_1_,*a*_2_),*b*); *G*_2_ = ((*a*_2_, *b*), *a*_1_); and *G*_3_ = ((*b, a*_1_), *a*_2_). If the first coalescence is more recent than population divergence (with *t*_1_ < *τ*), the gene trees are labelled *G*_1*a*_, *G*_2*a*_, *G*_3*a*_; otherwise they are labelled *G*_1*b*_, *G*_2*b*_, *G*_3*b*_. The *gdi* is the probability that the two *A* sequences coalesce first and before the population split: *gdi* = ℙ (*G*_1*a*_). Note that if there is no gene flow between *A* and *B*, gene trees *G*_2*b*_ and *G*_3*b*_ are impossible.

*P*_1_ is a simple function of 2*τ*_*AB*_/*θ* _*A*_ = *T*_*AB*_ (2*N*_*A*_), which is known as branch length in coalescent units (with one coalescent time unit to be 2*N*_*A*_ generations in population *A*). As *P*_1_ ranges from 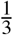(when populations *A* and *B* are at panmixia) to 1 (when *A* and *B* are completely isolated), Jackson *et al*. (2017) rescaled it so that the resulting *gdi* ranges from 0 to 1:

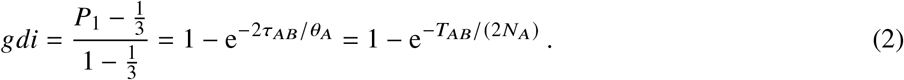

A *gdi* close to 1 indicates a high level of population divergence. Based on a meta-analysis of data from Pinho and Hey (2010), Jackson *et al*. (2017) suggest that populations are likely to be a single species if *gdi* < 0.2, and separate species if *gdi* > 0.7. Intermediate values (0.2 < *gdi* < 0.7) indicate ambiguous species status judged by the criterion.

Leaché *et al*. (2019) described a hierarchical merge algorithm for species delimitation based on *gdi*. Given a set of populations and a guide tree for them, the procedure attempts to merge two populations into one species, judged by *gdi*. Here we develop a python pipeline to automate the procedure. We include a hierarchical split algorithm as well. The hierarchical procedure of Leaché *et al*. (2019) relied on the MSC model without gene flow. In our pipeline we account for gene flow by using the MSC-M model implemented in bpp (Flouri *et al*., 2023). We first describe the definition and computation of *gdi* when there is gene flow in the model. Then we discuss our new pipeline and illustrate it using a simulated dataset. We apply the pipeline to three empirical datasets, for giraffes, milksnakes, and sunfish.

## Theory and Methods

### Definition and computation of gdi under the migration model

Consider two *A* sequences and one *B* sequence at a locus. Under the MSC-M model for two populations (fig. 2a), the probability for gene tree *G*_1_ = ((*a*_1_, *a*_2_), *b*) depends on the model parameters:

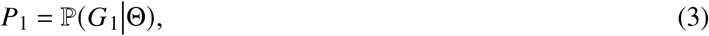

where Θ = (*τ*_*AB*_, *θ* _*A*_, *θ*_*B*_, *θ* _*AB*_, *M*_*AB*_, *M*_*BA*_) is the parameter vector. Jackson *et al*. (2017) estimated the minimum and maximum values for *P*_1_ to rescale *P*_1_ so that *gdi* falls into (0, 1). However, those limits depend on the model parameters. Here we redefine *gdi* as the probability that the first coalescence is between the two *A* sequences and that it occurs before reaching population divergence when we trace the genealogy of the three sequences backwards in time. In other words,

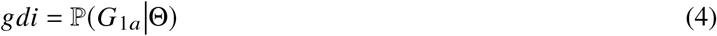

(fig. 1). In the special case of no gene flow, eq. 4 simplifies to eq. 2. There is no need for rescaling as 0 < ℙ (*G*_1*a*_)< 1.

**Figure 2:**
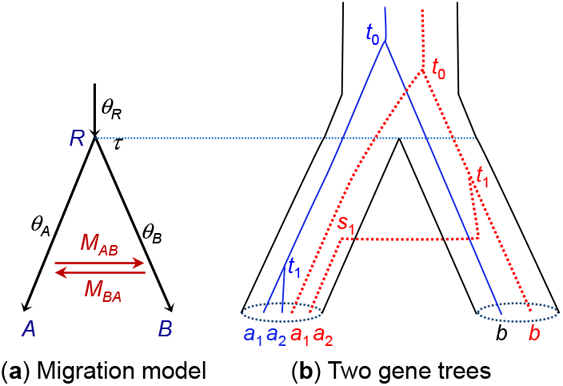
(**a**) An MSC-with-migration (MSC-M) model for two species or populations (*A, B*) showing the parameters. The two populations diverged time *τ ≡ τ*_*AB*_ ago and have since been exchanging migrants at the rate of *M*_*AB*_ = *m* _*AB*_*N*_*B*_ migrants per generation from *A* to *B* and at the rate *M*_*BA*_ = *m*_*BA*_*N*_*A*_ from *B* to *A*. (**b**) Two gene trees, each for two *A* sequences and one *B* sequence (*a*_1_, *a*_2_, *b*). In the blue tree (solid lines), *a*_1_ and *a*_2_ coalesce first, in population *A*, resulting in the gene tree *G*_1_ = ((*a*_1_, *a*_2_), *b*) (this is *G*_1*a*_ of fig. 1). In the red tree (dotted lines), *a*_2_ migrates into *B* and coalesce with *b* in *B*, resulting in the gene tree *G*_2_ = ((*a*_2_, *b*), *a*_1_) (this is *G*_2*a*_ of fig. 1).

Here we describe the computation of the *gdi* under the MSC-M model, following Leaché *et al*. (2019). Given two populations (*A* and *B*) with gene flow, the process of coalescent and migration when one traces the genealogical history of the sample of three sequences (*a*_1_, *a*_2_, and *b*) backwards in time can be described by a Markov chain, in which the states are specified by the number of sequences remaining in the sample and the population IDs (*A* and *B*) and sequence IDs (*a*_1_, *a*_2_, *b*) (table S1) (Hobolth *et al*., 2011; Zhu and Yang, 2012; Jiao and Yang, 2021). The initial state is 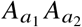 *B*_*b*_, with three sequences *a*_1_, *a*_2_, *b* in populations *A, A*, and *B*, respectively. This is also written ‘*AAB*’. State 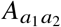 *B*_*b*_, abbreviated ‘*AB*_*b*_’, means that sequences *a*_1_ and *a*_2_ have already coalesced so that two sequences remain in the sample, with the ancestor of *a*_1_ and *a*_2_ in *A* while *b* is in *B*. Finally state *A*|*B* is an artificial absorbing state, in which all three sequences have coalesced with the sole ancestral sequence in either *A* or *B*. There are 21 states in the Markov chain, with the transition rate (generator) matrix Q = *q*_*ij*_ given in table S1.

The transition probability matrix over time *t* is then *P t* = {*p*_*ij*_ (*t*)} = e^*Qt*^, where *p*_*ij*_(*t*) is the probability that the Markov chain is in state *j* at time *t* (in the past) given that it is in state *i* at time 0 (the present time). Suppose *Q* has the spectral decomposition

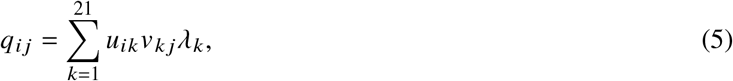

where 0 = *λ*_1_ > *λ*_2_ ≥ … ≥ *λ*_21_ are the eigenvalues of *Q*, and columns in *U* = {*u*_*ij*_} are the corresponding right eigenvectors, with *V* = {ν_*ij*_} = *U*^*−*1^. Then

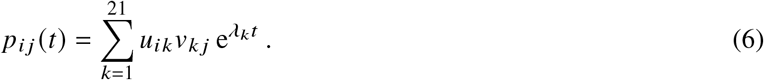

Gene tree *G*_1*a*_ arises if sequences *a*_1_ and *a*_2_ coalesce first and before *τ* (as in the blue gene tree of figure 2b). The coalescence can occur in either populations *A* or *B*. The coalescent time *t* has the density

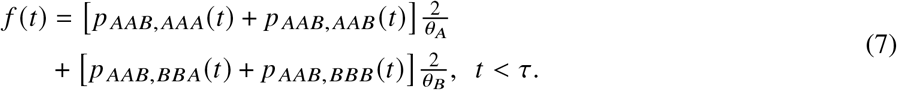

The two terms in the sum correspond to the coalescence occurring in *A* and *B*, respectively. For example, the first term is the probability that both *a*_1_ and *a*_2_ are in *A* right before time *t* (corresponding to states *AAA* or *AAB*), *p* _*AAB,AAA*_ (*t*) +*p* _*AAB,AAB*_ (*t*), times the coalescent rate. 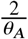 Similarly the second term is the probability density that *a*_1_ and *a*_2_ coalesce at time *t* in *B*, given by the probability that *a*_1_ and *a*_2_ are in *B* right before time *t* times the coalescent rate 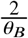

By averaging over the distribution of *t*, we have

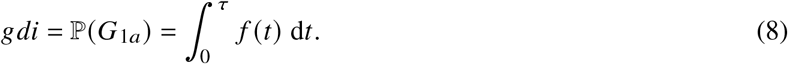

To calculate the integral in eq. 8, note that from eq. 6,

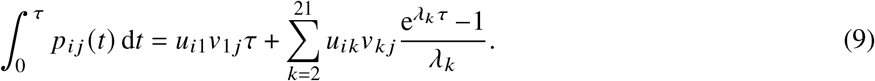

We have implemented this calculation of *gdi* in the python pipeline for the case where the two populations are sister lineages exchanging migrants between themselves but not with other populations.

When populations *A* and *B* are involved in gene flow with other populations, analytical calculation of the *gdi* becomes complicated. It is simpler to simulate gene trees for sequences *a*_1_, *a*_2_, *b* under the migration model involving all populations. Specifically, given the fully specified MSC-M model for all species/populations (including the species tree topology and parameters such as *τ, θ, M*), simulate gene trees with branch lengths (coalescent times) for a large number of loci (*R* = 10^6^, say), at which three sequences (*a*_1_, *a*_2_, *b*) are sampled. The *gdi* is simply the proportion of loci at which the gene tree is *G*_1*a*_, that is, *G*_1_ with *t*_1_ < *τ*_*AB*_ (figs. 1&2).

### The hierarchical merge and split algorithms

We implement both the hierarchical merge and hierarchical split algorithms in a python pipeline (fig. 3). Both algorithms require a guide tree for populations. Migration events involving both extant species/populations and extinct ancestral species/populations are allowed. In the merge algorithm, we progressively merge the populations into the same species, starting from the tips of the tree and moving towards the root. The merge is accepted if and only if the *gdi* < 0.2 for the population pair. The algorithm stops when no population pair can be merged (fig. 3a).

**Figure 3:**
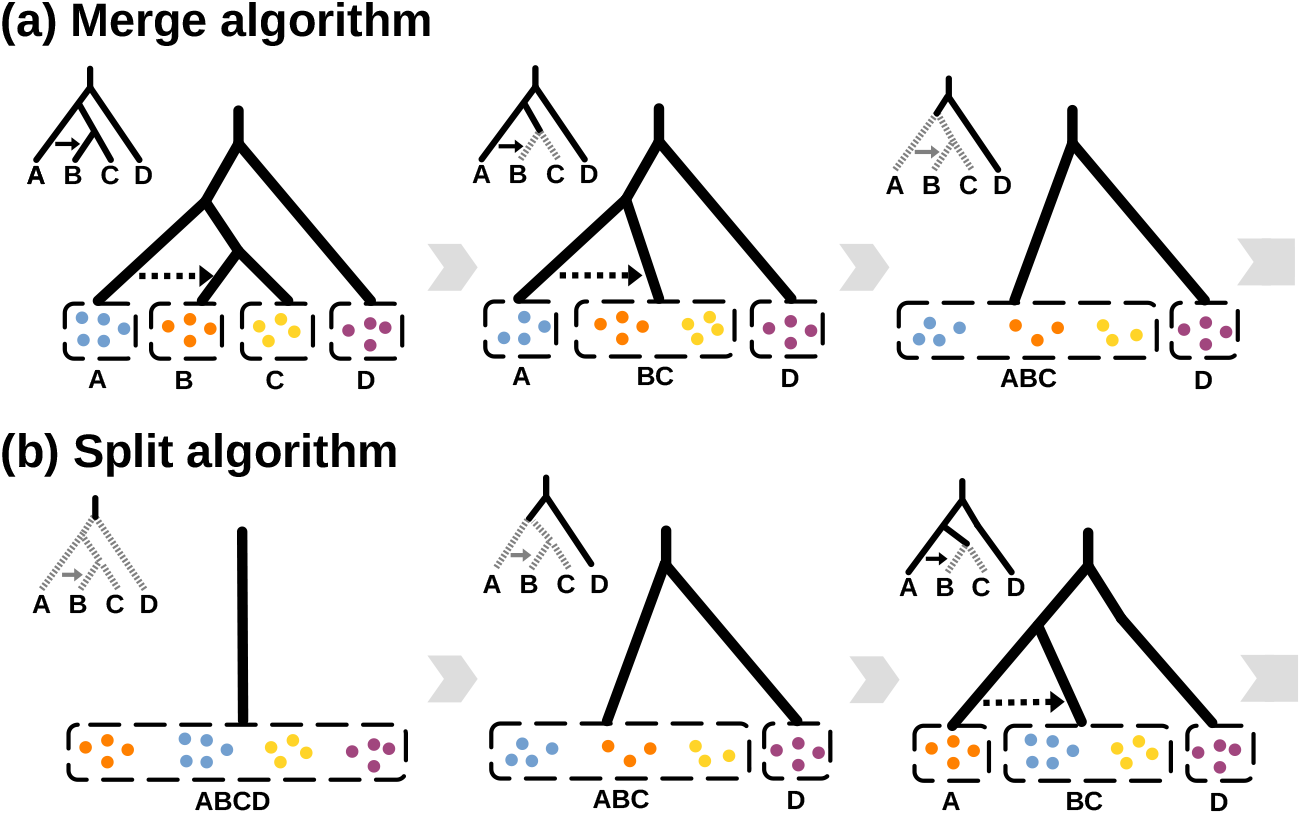
(**a**) Hierarchical merge and (**b**) hierarchical split algorithms applied to the same guide tree for four populations.

In the hierarchical split algorithm, we start from the MSC model of one species and progressively split each species into distinct species, starting from the root and moving towards the tips of the tree (fig. 3b). The split is accepted if and only if the *gdi* > 0.7. The algorithm stops when no species can be split (fig. 3b).

Both algorithms are implemented under either the MSC model with no gene flow (Rannala and Yang, 2003; Flouri *et al*., 2018) or the MSC-M model with continuous migration (Flouri *et al*., 2023). Under the MSC-M model, we retain the migration event when at least one of the two merged populations is involved in migration with a third species. For example, in the guide tree (initial delimitation) of figure 3a, there is migration from *A* to *B*. When *B* and *C* are merged into one species/population (*BC*), we retain the migration event (now from population *A* to population *BC*).

### Implementation of the hierarchical merge and split algorithms

Our new pipeline creates control files and Imap files to drive the analyses using bpp (an Imap file maps individual samples to species/populations under the specified species-delimitation hypothesis). It then examines the bpp output to decide on the next iteration of the hierarchical algorithm, generating new control and Imap files. The pipeline is itself driven by a control file. Many of the control variables are the same as used in bpp, and the same syntax is used between the two programs as much as possible.

Here we illustrate our pipeline through a re-analysis of the multilocus sequence data simulated under the isolation-by-distance model of figure 4a by Leaché *et al*. (2019). The control file is shown in figure 5. There are five populations, with *A, B, C, D* representing geographical populations of a species with a wide geographic distribution, while *X* is a new species that split off from population *A*. There is extensive gene flow between any two neighbouring populations of species *ABCD*, with migration rate *M* = *Nm* = 2 immigrants per generation, whereas there is no gene flow involving *X*. The data consisted of *L* = 100 simulated loci, with two diploid sequences sampled per species per locus, and 500 sites in the sequence.

**Figure 4:**
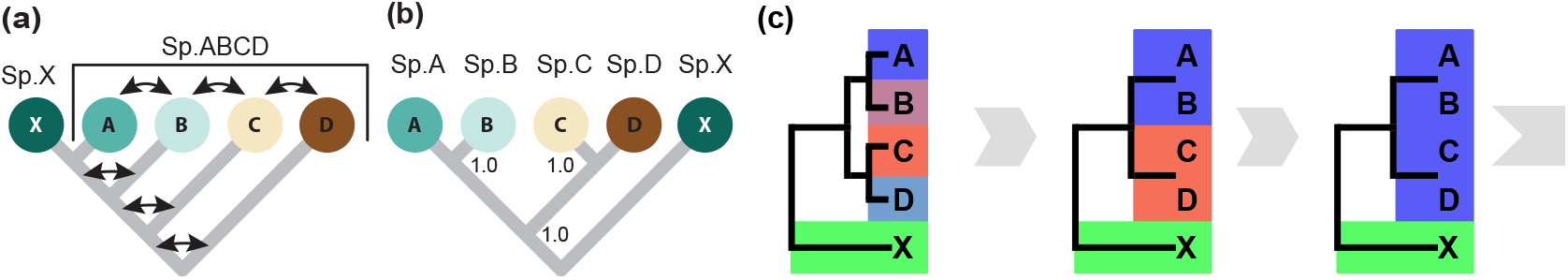
(**a**) An isolation-by-distance model used to simulate multilocus sequence data. The population divergence times (*τ*) are *τ*_*XABCD*_ = 0.04, *τ*_*XABC*_ = 0.03, *τ*_*XAB*_ = 0.02, and *τ*_*XA*_ = 0.01. The population size parameter is *θ* = 0.01 for all populations. The migration rate is *M* = *Nm* = 2 for any pair of adjacent populations in the species tree, and there is no gene flow involving the new species *X*. We simulated 100 loci, with 2 diploid sequences sampled per species per locus and 500 sites in the sequence. Redrawn after Leaché *et al*. (2019, fig. 5). (**b**) Incorrect species delimitation and phylogeny produced in Bayesian model selection using bpp under the MSC model assuming no gene flow, with every node receiving 100% posterior support. (**c**) Output from the pipeline providing a visual illustration of the progress of the hierarchical merge algorithm applied to the simulated data (see fig. 5 for the control file). The guide tree, (((*A, B*), (*C, D*), *X*), was inferred from bpp species tree estimation under the MSC model with no gene flow. The final delimitation from the algorithm has two species: *ABCD* and *X*.

**Figure 5:**
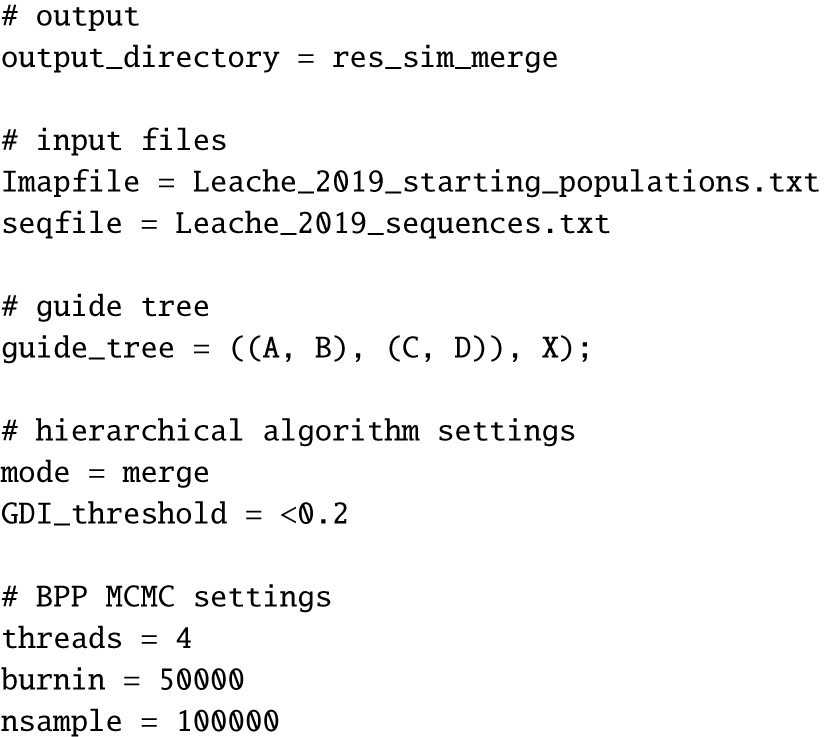
Control file for the merge analysis of the data simulated under the isolation-by-distance model of figure 4a. `output_directory` specifies the output directory in which result files will be written; `seqfile` is the sequence alignment file in phylip format; `Imapfile` specifies the mapping of individuals to populations; `guide_tree` is a Newick representation of the guide tree topology; and `mode` specifies the algorithm (`merge` or `split`). `GDI_threshold` specifies the *gdi* value below which two populations are merged into a candidate species. `threads` specifies the number of CPU threads used to run bpp, while `burnin`, `nsample` and `sampfreq` specify the MCMC settings for running bpp.

The guide tree (which is the starting delimitation for the merge algorithm) was generated using species tree estimation under the MSC model with no gene flow (i.e., the A01 analysis of Yang, 2015). The pipeline provides feedbacks about the current species delimitation and the decisions made during each iteration of the algorithm (figs. 4c&S1). In the first iteration, attempt was made to merge the two modern populations (*A* and *B*, and *C* and *D*). As *gdi* < 0.2 for each pair, both merges were accepted. In the second iteration, a merge between the pair *AB* and *CD* was attempted, and again this was accepted. In the third iteration, a merge between the pair *ABCD* and *X* was attempted. As *gdi* > 0.2, the merge was rejected. The final delimitation had two species, *ABCD* and *X*.

## Results

### *Species delimitation of giraffes (genus* Giraffa

Throughout the history of scientific studies of giraffes, their taxonomic position and classification has been controversial (Mitchell, 2009). Previous studies using morphological characters and molecular data produced inconsistent results, delimiting from one to six species in the *Giraffa* genus. Currently nine subspecies are recognised: *camelopardalis, angolensis, antiquorum, giraffa, peralta, reticulata, rothschildi, thornicrofti* and *tippelskirchi*. Petzold and Hassanin (2020) compiled a multilocus dataset of 21 introns (average sequence length 808 bp), sampled from 66 individuals from the nine subspecies, and conducted a number of population genetic and phylogenetic analyses. The authors suggested a delimitation with three species, although they noted that Bayesian model selection by bpp supported as many as five species.

We re-analyzed these data using our new pipeline, using the five-species phylogeny (fig. 6b) as the guide tree, which is also the starting delimitation in the merge algorithm. Based on phylogenetic analysis of mitochondrial haplotypes and identified hybrids (Fennessy *et al*., 2016; Petzold and Hassanin, 2020), bidirectional migration was specified between *reticulata* and the *tippelskirchi+thornicrofti* lineage, and between *reticulata* and the *camelopardalis+rothschildi+antiquorum* lineage. Migration rates were assigned the gamma prior G(0.1, 10) with mean 0.1/100 = 0.01 migrant individuals per generation. Merge and split analyses were conducted with the animal-specific *gdi* thresholds of 0.3 and 0.7, as recommended by Jackson *et al*. (2017) (see fig. S2 for the control file).

**Figure 6:**
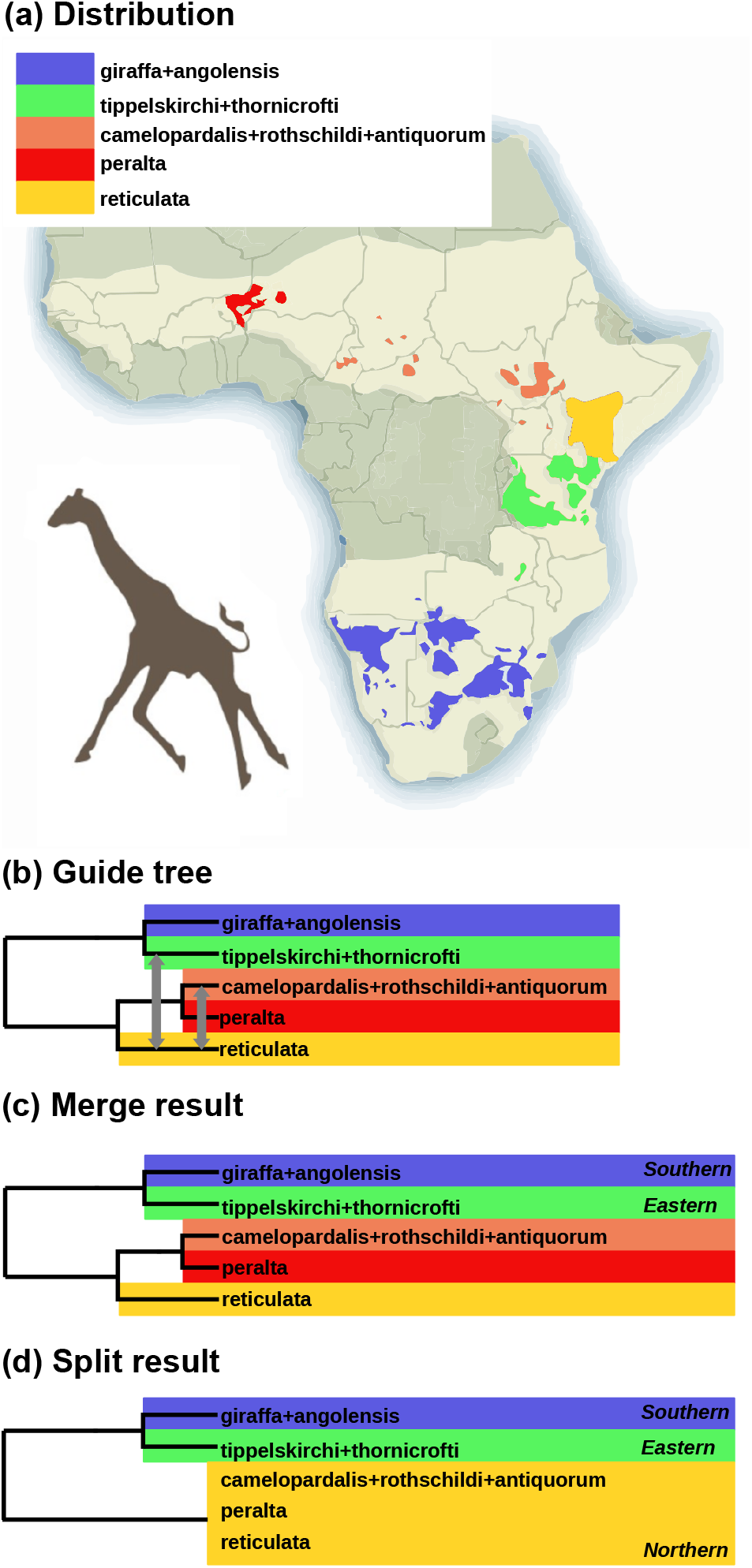
(**a**) Geographical distributions of five putative species within *Giraffa*. Bright region on map shows historical (ca. 1700) giraffe ranges (modified from https://giraffeconservation.org/giraffe-species/). (**b**) The guide tree for five populations of giraffes, with dotted lines indicating bidirectional migration events (Petzold and Hassanin, 2020, fig. 1). (**c**) The merge algorithm supports five species, while (**d**) the split algorithm supports three.

The merge algorithm suggested five species while the split algorithm suggested three (fig. 6c&d). Both methods recognized the Eastern (*tippelskirchi* and *thornicrofti*) and Southern (*giraffa* and *angolensis*) species present in the starting delimitation as distinct species. For the remaining Northern populations, the merge algorithm recognized three distinct species, while the split algorithm recognized only one.

Estimates of migration rates during the merge algorithm support the hypothesized patterns of gene flow between reticulated giraffes and the neighbouring populations (table 1). The highest migration rate was within the Northern populations from *cam*.*+rot*.*+ant*. to *reticulata*, which affected four of the 21 introns in the dataset according to Petzold and Hassanin (2020).

**Table 1.** Estimates (posterior means and 95% HPD CIs) of migration rates (*M* = *Nm*) between the five putative giraffe species in the guide tree of figure 6b.

| Donor | Recipient | $M$ (95% HPD CI) |
| --- | --- | --- |
| TipTho | reticulata | 0.002 (0.000, 0.015) |
| reticulata | TipTho | 0.002 (0.000, 0.009) |
| reticulata | CamRotAnt | 0.027 (0.000, 0.129) |
| CamRotAnt | reticulata | 0.123 (0.000, 0.328) |

### *Species delimitation in milksnakes (*Lampropeltis triangulum)

The American milksnake *Lampropeltis triangulum* is a New World snake with one of the widest known geographic distributions within the squamates. Seven subspecies are known: *abnorma, polyzona, micropholis, triangulum, gentilis, annulata*, and *elapsoides* (fig. 7a). Ruane *et al*. (2014) analyzed 11 nuclear loci (average length 537 bp) for 164 individuals from the seven subspecies using bpp model comparison and found evidence for seven distinct species. Chambers and Hillis (2020) re-analyzed these data and suggested that several species hypothesized by Ruane *et al*. (2014) may represent arbitrary slices of continuous geographic clines. They instead suggested two delimitation hypotheses, with three and one species, respectively, as shown in figure 7c&d.

**Figure 7:**
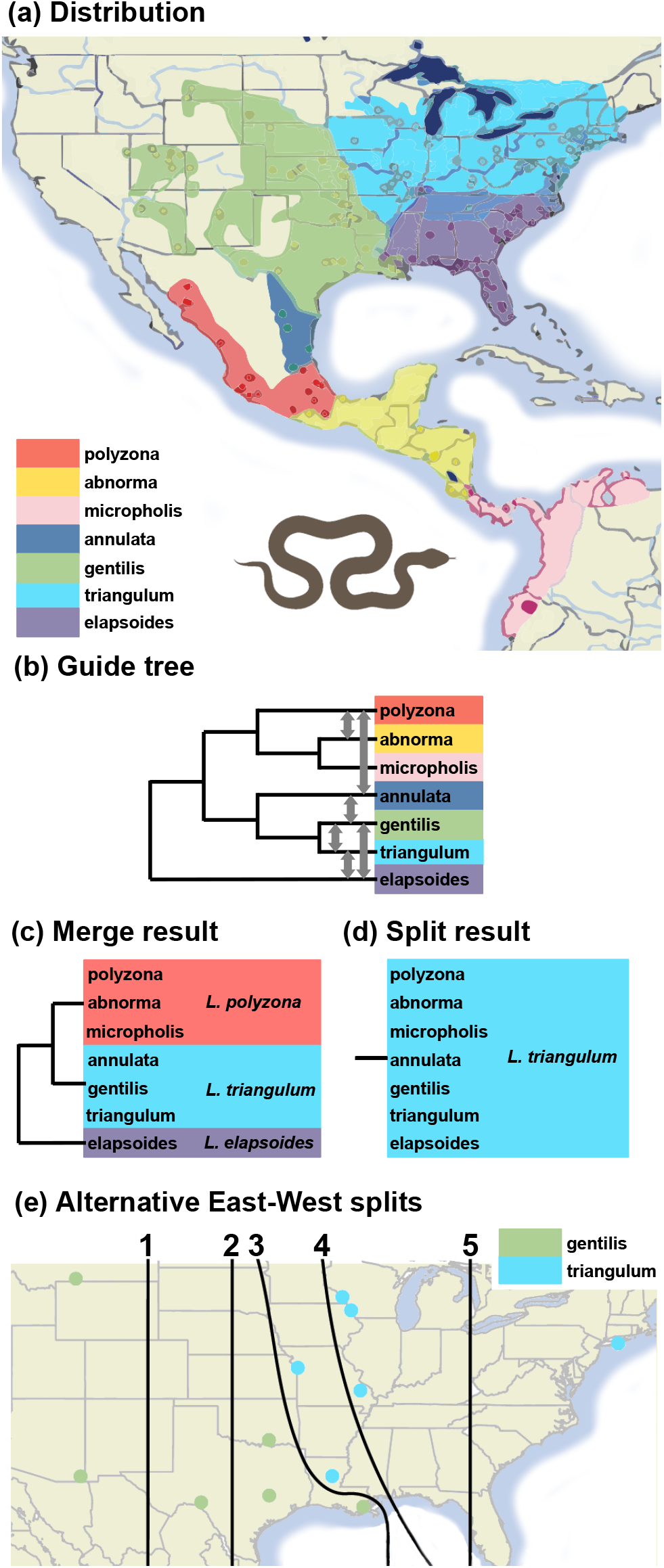
(**a**) Geographic distribution of seven milksnake subspecies (map based on and modified from Ruane *et al*. 2014, fig. 1d). (**b**) The guide tree with bidirectional migration events. (**c**&**d**) Inferred delimitation hypotheses by the merge and split algorithms. (**e**) Alternative delimitation hypotheses tested by Chambers and Hillis (2020), each of which splits the *gentilis* and *triangulum* samples at an arbitrary West-East divide line.

We reanalyzed the data of Ruane *et al*. (2014) using our pipeline, using the guide tree for seven populations of Chambers and Hillis (2020) (fig. 7b). As the original Ruane *et al*. (2014) analysis found ongoing genetic exchange between geographically adjacent populations, we added bidirectional migration events in the guide tree (fig. 7b). Merge and split algorithms were run using *gdi* thresholds of 0.3 and 0.7 (see fig. S3 for control file).

The merge algorithm suggested a maximum of three species, grouping the subspecies *abnorma, polyzona*, and *micropholis* into one species, and *triangulum, gentilis*, and *annulata* into another species (fig. 7c). This is the same delimitation as the three-species hypothesis of Chambers and Hillis (2020). The split analysis supported only one species (fig. 7d).

Migration rates between the adjacent subspecies/populations during the merge analysis suggested ongoing genetic exchange between some of the subspecies pairs, in particular, between *L. annulata* and *L. gentilis* and between *L. abnorma* and *L. polyzona* (table 2).

**Table 2.** Estimates of migration rates (*M*) between seven milksnake populations during the merge algorithm (fig. 7)

| It. | Donor | Recipient | *M* (95% HPD CI) |
| --- | --- | --- | --- |
| 1 | elapsoides | gentilis | 0.003 (0.000, 0.019) |
|  | gentilis | elapsoides | 0.010 (0.000, 0.055) |
|  | elapsoides | triangulum | 0.009 (0.000, 0.052) |
|  | triangulum | elapsoides | 0.055 (0.000, 0.152) |
|  | annulata | polyzona | 0.002 (0.000, 0.012) |
|  | polyzona | annulata | 0.003 (0.000, 0.016) |
|  | annulata | gentilis | 0.162 (0.000, 0.331) |
|  | gentilis | annulata | 0.053 (0.000, 0.220) |
|  | polyzona | abnorma | 0.044 (0.000, 0.189) |
|  | abnorma | polyzona | 0.127 (0.000, 0.293) |
|  | gentilis | triangulum | 0.011 (0.000, 0.070) |
|  | triangulum | gentilis | 0.050 (0.000, 0.267) |
| 2 | elapsoides | GenTri | 0.008 (0.000, 0.049) |
|  | GenTri | elapsoides | 0.067 (0.000, 0.142) |
|  | annulata | PolAbn | 0.002 (0.000, 0.010) |
|  | PolAbn | annulata | 0.003 (0.000, 0.020) |
|  | annulata | GenTri | 0.081 (0.000, 0.276) |
|  | GenTri | annulata | 0.073 (0.000, 0.181) |
| 3 | elapsoides | AnnGenTri | 0.016 (0.000, 0.085) |
|  | AnnGenTri | elapsoides | 0.102 (0.028, 0.184) |
|  | MicPolAbn. | AnnGenTri | 0.006 (0.000, 0.034) |
|  | AnnGenTri | MicPolAbn | 0.080 (0.028, 0.135) |

Chambers and Hillis (2020) also applied an arbitrary West-East divide to split the *gentilis* and *triangulum* populations into two species, thus generating five arbitrary delimitation hypotheses (each with two species) (fig. 7e). They found that all five delimitation hypotheses were supported by Bayesian model selection using bpp, even though they are not mutually compatible.

We used our new pipeline to conduct a re-analysis to test the five delimitation hypotheses, using the merge algorithm with the same settings as above. The data consisted of only the 38 individuals from *gentilis, triangulum*, and *annulata* populations. The same guide tree for the three populations is used, but each hypothesis is represented by constructing an Imap file to map the individual samples to the three populations (see figs. S4&S5 for the control file and command-line scripts). Bidirectional migration between *gentilis* and *triangulum* is allowed in the guide tree.

Under each of the five delimitation hypotheses, the merge algorithm merged the two subspecies *gentilis* and *triangulum* into a single species.

### *Introgression and species delimitation in the longear sunfish (*Lepomis megalotis)

The longear sunfish (*Lepomis megalotis*) is a freshwater fish in the sunfish family, Centrarchidae, of order Perciformes. It is native to eastern North America from the Great Lakes down to northeastern Mexico (fig. 8a). Six subspecies are recognised: *aquilensis, solis, ouachita, megalotis, ozark*, and *pelastes*. Due to widespread geographic distribution and frequent hybridization, species delimitation in the longear sunfish poses considerable challenges.

**Figure 8:**
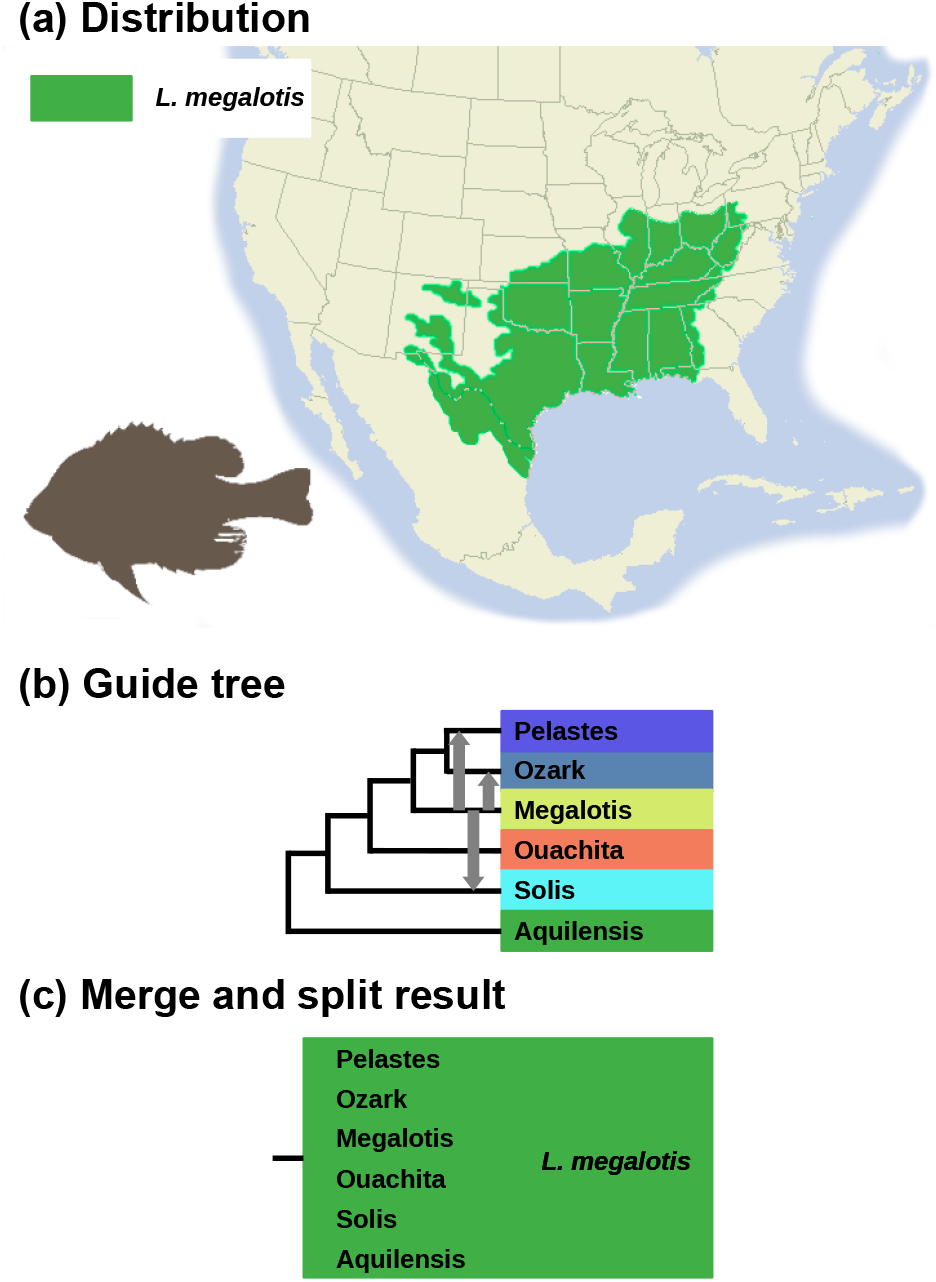
(**a**) Geographic distribution of longear sunfish (*Lepomis megalotis*) (map based on http://www.roughfish.com/content/longear-sunfish). (**b**) The guide tree, with three migration events (from *L. megalotis* to *L. pelastes, L. solis*, and *L. ozark*) indicated by arrows. (**c**) Both merge and split algorithms support a single species.

Kim *et al*. (2022) analyzed a dataset of 163 ddRAD loci (average sequence length 89 bp) sampled from 50 individuals from the six subspecies. After determining a species/population phylogeny using IQ-tree, they analyzed the data under the MSC model with no gene flow using bpp to calculate the *gdi* scores, to delimit species in the group. They found that none of the populations had *gdi* values supporting distinct species status. Kim *et al*. (2022) also found evidence for multiple instances of historical or ongoing gene flow.

We reanalyzed the data of Kim *et al*. (2022), using the MSC-M model to calculate *gdi*, accommodating migration between the subspecies. Based on the hybridization patterns observed by Kim *et al*. (2022), migration from *megalotis* to *pelastes, solis*, and *ozark* was specified in the guide tree (fig. 8b). Migration rates were assigned the gamma prior G(0.1, 10) with mean 0.01 migrant individuals per generation. Merge and split algorithms were run using *gdi* thresholds of 0.3 and 0.7 (control file in fig. S6).

Both merge and split analyses supported a single species. This is congruent with the delimitation of Kim *et al*. (2022), in which *gdi* was calculated under the MSC model without gene flow. Estimates of the migration rates between the subspecies during the merge algorithm (table 3) were consistently large, supporting the classification of those populations as a single species.

**Table 3.** Estimates of migration rates between five sunfish populations during the merge algorithm (fig. 8)

| It. | Donor | Recipient | *M* (95% HPD CI) |
| --- | --- | --- | --- |
| 1 | megalotis | solis | 0.605 (0.412, 0.808) |
|  | megalotis | pelastes | 0.537 (0.370, 0.709) |
|  | megalotis | ozark | 0.322 (0.103, 0.596) |
| 2 | megalotis | solis | 0.692 (0.462, 0.945) |
|  | megalotis | PelOzk | 0.693 (0.397, 0.989) |
| 3 | PelOzkMeg | Solis | 0.579 (0.387, 0.785) |
| 4 | PelOzkMegOua | Solis | 0.407 (0.279, 0.541) |

## Discussion

### The guide tree, gene flow, and paraphyletic species

Our pipeline requires the user to supply a guide tree. This may be inferred using a species tree estimation method under the MSC model with no gene flow (Yang and Rannala, 2014; Rannala and Yang, 2017). Alternatively phylogenetic programs such as IQ-tree (Minh *et al*., 2020) and RAxML (Stamatakis *et al*., 2012) can be used to infer the maximum likelihood tree using concatenated genomic data or mitochondrial genomic sequences.

We note that the hierarchical merge and split algorithms implicitly assume a monophyletic species definition and thus do not work when a species is paraphyletic. Paraphyletic species or species comprising of multiple populations that are not monophyletic are common (Crisp and Chandler, 1996). The model tree of figure 4a represents such a scenario, in which species *ABCD* is paraphyletic. The issue here concerns the non-monophyly of the populations of the same species, and is different from the monophyly of a gene tree, which is problematic if used as a criterion for species delimitation (Knowles and Carstens, 2007). In other words, non-monophyly of gene trees is a natural consequence of the coalescent process under the MSC model and can arise even if each species on the species tree is monophyletic. We note that the reversible-jump algorithm for Bayesian model comparison of Yang and Rannala (2010) is similarly based on the assumption that species should be monophyletic and has the same problem. Crisp and Chandler (1996) argued that while allowing for paraphyletic species, one may still insist on higher taxa being always monophyletic.

If all populations are completely isolated with no gene flow, the concept of a paraphyletic species does not appear very sensible. For example, if the population phylogeny is the model of figure 4a but without gene flow, i.e., ((((*X, A*), *B*), *C*), *D*), it appears nonsensical to designate population *X* as a distinct species from *ABCD* while lumping populations *A, B, C*, and *D* into one species, given that populations *B, C*, and *D* split from *A* earlier than *X* did. However, with gene flow between populations, the population divergence history may render the species to be paraphyletic (as in the model of fig. 4a with gene flow). Here we discuss three possible heuristic-delimitation strategies to accommodate paraphyletic species.

The first is to use a guide tree for all populations (including those that make up the paraphyletic species) assuming no gene flow. This is used in Leaché *et al*. (2019) and in this paper (fig. 4c), where the guide tree is constructed under the MSC model ignoring gene flow and then used to calculate the *gdi*. The resulting guide tree (fig. 4c; see also Leaché *et al*., 2019, fig. 3b) may reflect gene flow and differ from the history of species/population divergence. For the example dataset of figure 4a, this approach led to the correct delimitation of two species (*ABCD* and *X*) even though the guide tree did not have the correct topology.

The second strategy is to use the MSC-M model accommodating gene flow in the guide tree (e.g., the species tree or MSC-M model of fig. 4a), but allow the merge of nonsister lineages (e.g., *A* and *B*, or *B* and *C*, or *C* and *D* in the guide tree of fig. 4a) during the hierarchical merge algorithm if there is significant evidence for gene flow between them and if the rate of gene flow exceeds a certain cutoff. Thus two populations may be merged into one species either because their divergence is low (*gdi* < 0.2) or because the migration rate is high (e.g., *M* = *Nm* > 0.1). When two nonsister lineages on the guide tree are merged, one may use the idea of *displayed species trees* (Degnan, 2018) to generate the new species tree or model. For example, if the migration rate *M*_*BD*_ exceeds a cutoff with significant gene flow from *B* to *D*, population *D* will be merged into *B* so that the species tree becomes (((*X, A*), (*B, D*)), *C*), whilst if *M*_*DB*_ is significant and exceeds the cutoff, population *B* is merged into *D* so that the species tree becomes (((*X, A*), *C*), (*B, D*)). We may allow multiple *gdi*-based merges but only one migration rate-based merge in each hierarchical merge step.

A third strategy may be to infer the history of population divergence and gene flow and leave it to the taxonomist to integrate the information with other sources of evidence to reach a decision on the species status of the populations. Suppose the analysis of the genomic data allows us to infer that the MSC-M model of figure 4a is a good representation of the history for the five populations, with parameter estimates close to the true values used to generate the data. The taxonomist may decide that there are two species, with *ABCD* to be a paraphyletic species.

We also note that where there is gene flow between species or populations, multiple strategies exist in the framework of Bayesian model selection as well. Given two populations (*A, B*), three models or scenarios may be considered: (i) *H*_1_: one single species, (ii) *H*_2ø_: two species with no gene flow, and (iii) *H*_2*m*_: two species with gene flow (with either *M*_*AB*_ > 0 or *M*_*BA*_ > 0 or both). Leaché *et al*. (2019) compared *H*_1_ and *H*_2ø_ to decide whether there is one or two species, and noted that if there is a population split followed by gene flow so that *H*_2*m*_ is the true model, then *H*_2ø_ wins over *H*_1_, potentially leading to false positives (over-splitting) by the approach of model selection. Alternatively one may insist on species status only if there is no significant amount of gene flow, that is, only if *H*_2ø_ wins over both *H*_1_ and *H*_2*m*_). This is arguably a more faithful implementation of the Biological Species Concept (Dobzhansky, 1937; Mayr, 1942; Coyne and Orr, 2004) than the comparison between *H*_1_ and *H*_2ø_ (Yang and Rannala, 2010), although it may suffer from high false negatives (over-lumping).

### Utility and challenges of heuristic species delimitation

In this paper we have developed a python pipeline to automate hierarchical merge and split algorithms for heuristic species delimitation. The merge algorithm was described and applied by Leaché *et al*. (2019), and here we have made the procedure automatic. We have also implemented the hierarchical split algorithm. Our tests using both simulated and empirical datasets suggest that the implementation is correct. Heuristic species delimitation based on criteria such as the *gdi* appear less likely to suffer from over-splitting, which has been discussed extensively as a problem with the approach of Bayesian model selection (Yang and Rannala, 2010).

While we have used in this paper the *gdi* as the criterion in the hierarchical merge and split algorithms, alternative heuristic criteria may be applied in our pipeline. It is also possible to apply a composite criterion. For example, we may insist, besides the *gdi* cutoff, that the species split time reach a minimum of 10^4^ generations (Rannala and Yang, 2020) and that the (effective) migration rate between two species should not exceed *M* = *Nm* = 0.1 migrants per generation. When there exist contact zones between populations and one can estimate the proportion of hybridization and back-crossing (*h*), an appropriate criterion may be the rate ratio *m*/*h*, which measures reproductive isolation: a value of 1 means that introgressed alleles are neutral and have the same chance of being retained as a native allele in the recipient population, while *m* / *h ≪* 1 means that introgressed alleles are strongly deleterious and purged from the population by natural selection, indicating the existence of (post-zygotic) reproductive isolation (Westram *et al*., 2022). Note that genetic/genomic data can be used to identify hybrids and estimate *h* (Anderson and Thompson, 2002; Chakraborty and Rannala, 2023) and to estimate the effective migration rate *m* (Hey, 2010; Hey *et al*., 2018; Gronau *et al*., 2011; Flouri *et al*., 2023). It may also be noted that a seemingly small migration rate (*m*), in the order of *M* = *Nm* = 0.5 may have a major impact on the genetic history of the populations (Long and Kubatko, 2018; Jiao and Yang, 2021).

Here we note a few limitations of *gdi* and our pipeline, which reflect challenges for heuristic delimitation in general. First, the *gdi* and thus our pipeline may suffer from ambiguities. Suppose there are *K* populations on the guide tree, the merge algorithm may arrive at a high number of species (*K*_*u*_) while the split algorithm at a low number (*K*_*l*_), with 1*≤ K*_*l*_ *≤ K*_*u*_ *≤ K*. When the two algorithms disagree (with *K*_*l*_ < *K*_*u*_), the *gdi* is ambiguous (Jackson *et al*., 2017). For example, this occurred in both the giraffe and milksnake datasets (figs. 6c&d and 7c&d), although both algorithms suggested one species in the sunfish dataset (fig. 8c). If taxonomists may be classified into “splitters” and “lumpers”, a splitter may prefer the merge algorithm while a lumper may use the split algorithm.

Another ambiguity concerns the definition of *gdi*. In eqs. 2 or 4, we considered a sample of two *A* sequences and one *B* sequence. One may similarly consider two *B* sequences and one *A* sequence. Under the MSC model with no gene flow, the two definitions are

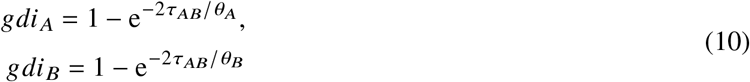

(cf: eq. 2). When *A* and *B* have very different population sizes, the two definitions may be inconsistent concerning the species status of *A* and *B*. For example, if *N*_*A*_ *≪ N*_*B*_, population *A* may appear to be a distinct species from *B* judged by *gdi* _*A*_, but *B* may not appear to be a different species from *A* judged by *gdi*_*B*_ (Leaché *et al*., 2019). One may avoid such awkwardness by insisting on both *gdi* _*A*_ and *gdi*_*B*_ exceeding the threshold in the pipeline.

Second, it is easy to imagine scenarios in which the *gdi* or our pipeline may generate either false positives (the error of over-splitting) or false negatives (the error of over-lumping). For example, a large *gdi* may result from a very small population size even if the divergence between the populations is recent (i.e., small *θ* and small *τ*), leading to false positives. Thus it may be advisable to apply an absolute divergence time threshold together with the *gdi* (Rannala and Yang, 2020). Similarly *gdi* may suffer from false negatives, especially in case of recent radiative speciations where not enough time has passed to generate sufficient genetic divergence between species throughout the genome (as in the ciclid fishes from the African lakes, Malinsky *et al*., 2018).

The greatest challenge to heuristic species delimitation, when applied to allopatric geographic populations, may be the arbitrary nature of the species concept (e.g., De Queiroz, 2007; Mallet *et al*., 2023). For example, Darwin considered the difference between a species and a variety (population, race, or subspecies) to be one of degree, while Bateson (1909) considered species to be real and suggested hybrid sterility as a test of species status. The latter view is epitomized in the Biological Species Concept Dobzhansky (1937); Mayr (1942); Coyne and Orr (2004), which emphasizes reproductive isolation.

Note that heuristic species delimitation discussed in this paper has a lot of similarities to early heuristics including genetic-distance cutoffs (such as the ‘10x rule’ in DNA bar-coding, Hebert *et al*., 2004) and reciprocal monophyly of gene trees (Baum and Shaw, 1995), and may be considered refined versions of those earlier criteria. For example, under the complete-isolation model (MSC with no gene flow), *P*_1_ and *gdi* (eqs. 1&2) are simple functions of *τ* / (*θ* / 2) = *T*/(2*N*), and both contrast within-species polymorphism with between-species divergence, just as the ‘10x rule’ does — note that 2*N* is the average divergence time (in generations) between two sequences sampled from the same species (of size *N*) while *T* is the species divergence time (in generations). Similarly gene tree *G*_1_ = ((*a*_1_, *a*_2_, *b*) is one of within-species monophyly given the three sequences at the locus (*a*_1_, *a*_2_, *b*). A major difference is that earlier criteria depend on simple features of the genetic data, whereas the methods discussed here are formulated using population parameters. From the viewpoint of statistical inference, the distinction between data and their summaries on one hand and model and parameters on the other is important, as it allows us to leverage the power of genomic data and the statistical theory that justifies the estimation method. For example, earlier methods may suffer from a lack of phylogenetic information at a locus when sequence alignments are used to infer gene trees, and incomplete lineage sorting or stochastic fluctuations of the coalescent process, while those concerns are addressed in methods discussed here. By analyzing multilocus genomic data under the MSC model, reliable estimation of the species tree and population parameters is possible even if all gene trees are poor (Xu and Yang, 2016).

Nevertheless, heuristic species delimitation discussed here suffer from the arbitrary nature of species definition, just as did the earlier heuristics, when applied to allopatric populations that do not overlap in their geographical distributions. Even if a full characterization of the history of the populations is available, in terms of the order and timings of population splits, population sizes, and the directions, timings and strengths of gene flow between populations, a universally accepted view on species status may not exist. Allopatric populations, with no or highly restricted gene flow between them, may be classified as distinct species, or merely subspecies or populations, and some arbitrariness appears unavoidable. We consider the role of the MSC model and the genomic data to be providing an accurate and detailed characterization of the history of population divergence and gene flow, as can be achieved through analyzing genomic data using powerful statistical methodologies such as Bayesian model selection and parameter estimation. In this regard, no fundamental difference is seen to exist between model comparison of Yang and Rannala (2010) and parameter estimation discussed here (Leaché *et al*., 2019), as both aim to characterize the history of population/species divergence and gene flow.

While acknowledging those caveats, we suggest that our pipeline allows one to utilize the power of the MSC framework and the bpp program to estimate population parameters precisely and accurately using the ever-increasing genomic sequence data. In particular, the recently implemented MSC-M model in bpp accommodates gene flow (continuous migration) between species, subspecies or populations (Flouri *et al*., 2023). We hope that our pipeline may become a useful tool for evolutionary biologists to assess and validate genomic-based species delimitation by integrating multiple lines of evidence, including morphological and behaviorial characteristics, and patterns of hybridization (Fujita *et al*., 2012; Solis-Lemus *et al*., 2015; Kim *et al*., 2022).

## Program Availability

The pipeline, called hhsd (for Hierarchical Heuristic Species Delimitation), is written in python, and drives parameter estimation under the MSC or MSC-M models using bpp. The source code, documentation, and empirical datasets analyzed in the paper are available at https://github.com/abacus-gene/hhsd.

## Supplementary Material

Data available from the Dryad Digital Repository: https://doi.org/10.5061/dryad.jm63xsjhc

## Acknowledgements

We thank Jim Mallet and Bruce Rannala for discussions, and Adam Leaché and Jim Mallet for many constructive comments and criticisms. We thank Asif Tamuri for reviewing the code.

