## Supplementary Figures for "Hierarchical heuristic species delimitation under the multispecies coalescent model with migration"

```
Number of species in starting delimitation:  5
((A, B), (C, D)), X);

*** Iteration 1 ***

Estimated tau and theta parameters:
        theta                   tau
X       0.0098 (0.0085, 0.0112)
A       0.0138 (0.0068, 0.0227)
B       0.0202 (0.0083, 0.0372)
C       0.0212 (0.0074, 0.0398)
D       0.0263 (0.0087, 0.0530)
ABCDX   0.0368 (0.0324, 0.0412) 0.0101 (0.0093, 0.0108)
ABCD    0.0432 (0.0394, 0.0458) 0.0006 (0.0005, 0.0008)
AB      0.0128 (0.0063, 0.0197) 0.0003 (0.0002, 0.0005)
CD      0.0204 (0.0085, 0.0324) 0.0002 (0.0001, 0.0005)

Proposal results:
Node pair        gdi 1   gdi 2   merge accepted?
'A', 'B'         0.05    0.03    True
'C', 'D'         0.03    0.02    True

Number of species after iteration 1:  3
((AB, CD), X);

*** Iteration 2 ***

Estimated tau and theta parameters:
        theta                   tau
X       0.0098 (0.0085, 0.0112)
AB      0.0194 (0.0146, 0.0243)
CD      0.0286 (0.0207, 0.0368)
ABCDX   0.0367 (0.0323, 0.0410) 0.0102 (0.0094, 0.0109)
ABCD    0.0428 (0.0395, 0.0466) 0.0006 (0.0005, 0.0009)

Proposal results:
Node pair        gdi 1   gdi 2   merge accepted?
'AB', 'CD'       0.07    0.05    True

Number of species after iteration 2:  2
(ABCD, X);

*** Iteration 3 ***

Estimated tau and theta parameters:
        theta                   tau
X       0.0098 (0.0086, 0.0112)
ABCD    0.0441 (0.0411, 0.0473)
ABCDX   0.0365 (0.0322, 0.0410) 0.0102 (0.0095, 0.0110)


Proposal results:
Node pair        gdi 1   gdi 2   merge accepted?
'ABCD', 'X'      0.39    0.87    False

Number of species after iteration 3:  2
(ABCD, X);

Final delimitation reached.
```

Figure S1: Screen output from running the hierarchical merge algorithm to analyze the simulated dataset of figure 4.

```
# Notes: species are renamed as follows
#
# gir_ang = giraffa+angolensis
# tip_tho = tippelskirchi+thornicrofti
# cam_rot_ant = camelopardalis+rothschildi+antiquorum
# per = peralta
# ret = reticulata

# output
output_directory = res_giraffe_merge

# input files
Imapfile = Imap_Giraffe.txt
seqfile = MSA_Giraffe.txt

# guide tree
guide_tree = ((gir_ang,tip_tho),((cam_rot_ant,per),ret));

# migration events and priors
migration = {
    ret <-> tip_tho,
    ret <-> cam_rot_ant,
}
migprior = 0.1 10

# hierarchical algorithm settings
mode = merge
gdi_threshold = <0.3

# BPP MCMC settings
threads = 8
burnin = 200000
nsample = 500000
```

Figure S2: Control file for the merge algorithm in analysis of the giraffe data. The control file for the split algorithm is the same except for the hierarchical-algorithm settings: `mode = merge` and `gdi_threshold = >0.7`. The results are in figure 6.

```
# Notes: species are renamed as follows
#
# Po = polyzona
# Ab = abnorma
# Mi = micropholis
# An = annulata
# Ge = gentilis
# Tr = triangulum
# El = elapsoides

# output
output_directory = res_milksnake_merge

# input files
Imapfile = Imap_Lampropeltis.txt
seqfile = MSA_Lampropeltis.txt

# guide tree
guide_tree = (((Mi, (Po, Ab)), (An, (Ge, Tr))), El);

# migration events and priors
migration = {
    Po <-> Ab,
    Po <-> An,
    An <-> Ge,
    Ge <-> Tr,
    Ge <-> El,
    Tr <-> El,
}
migprior = 0.1 10

# hierarchical algorithm settings
mode = merge
gdi_threshold = <0.3

# BPP MCMC settings
threads = 8
burnin = 200000
nsample = 500000
```

Figure S3: Control file for the merge analysis of the milksnakes data. The control file for the split algorithm is the same except for the hierarchical-algorithm settings: `mode = merge` and `gdi_threshold = >0.7`. The results are presented in figure 7.

```
# output
output_directory = # will be set from the command line

# input files
Imapfile = # will be set from the command line
seqfile = trigentalt.txt

# guide tree
guide_tree = ((Ge, Tr), Al);

# migration events and priors
migration = { Ge <-> Tr }
migprior = 0.1 10


# hierarchical algorithm settings
mode = merge
gdi_threshold = <0.3

# BPP MCMC settings
threads = 8
burnin = 200000
nsample = 500000
```

Figure S4: Control file for the analysis of the milksnake data under delimitation hypotheses reflecting the East-West splits (fig. 7e). The `Imapfile` and `output_directory` parameters are left empty, as they will be provided at the command line (fig. S5), to ensure that they correspond to the five delimitation hypotheses being tested.

```
hhsd --cfile cf_milksnake_EW.txt --cfpor \
Imapfile = 1alt.Imap.txt, output_directory = res_EW_1

hhsd --cfile cf_milksnake_EW.txt --cfpor \
Imapfile = 2alt.Imap.txt, output_directory = res_EW_2

hhsd --cfile cf_milksnake_EW.txt --cfpor \
Imapfile = 3alt.Imap.txt, output_directory = res_EW_3

hhsd --cfile cf_milksnake_EW.txt --cfpor \
Imapfile = 4alt.Imap.txt, output_directory = res_EW_4

hhsd --cfile cf_milksnake_EW.txt --cfpor \
Imapfile = 5alt.Imap.txt, output_directory = res_EW_5
```

Figure S5: Shell script used to iterate through the five East-West delimitation hypotheses for the milksnakes (fig. 7e). The `-cfpor` (control file parameter override) flag is used to override parameters of the control file via the command-line interface, setting the Imap file to match the delimitation hypothesis, and specifying the output directories for each analysis.

```
# Notes: species are renamed:
#
# PEL = pelastes
# OZK = ozark
# MEG = megalotis
# LIT = ouachita
# SOL = solis
# AQU = aquilensis

# output
output_directory = res_sunfish_merge

# input files
Imapfile =  Imap_Sunfish.txt
seqfile = MSA_Sunfish.txt

# guide tree
guide_tree = (((((PEL,OZK),MEG),LIT),SOL),AQU);

# migration events and priors
migration = {
    MEG -> PEL,
    MEG -> SOL,
    MEG -> OZK,
}
migprior = 0.1 10

# hierarchical algorithm settings
mode = merge
gdi_threshold = <0.3

# BPP MCMC settings
threads = 16
burnin = 200000
nsample = 500000
```

Figure S6: Control file for the merge analysis of the sunfish data (fig. 8). The control file for the split algorithm is the same except for the hierarchical-algorithm settings: `mode = merge` and `gdi_threshold = >0.7`.

Table S1: Rate matrix for Markov chain describing transitions between states in multispecies coalescent with migration model with two populations ($A$ and $B$) and three sequences ($a_1$, $a_2$, and $b$).

| | $AAA$ | $AAB$ | $ABA$ | $ABB$ | $BAA$ | $BAB$ | $BBA$ | $BBB$ | $A_{a_1}A$ | $A_{a_2}A$ | $A_bA$ | $C_{a_1}B$ | $B_{a_2}B$ | $B_bB$ | $A_{a_1}B$ | $A_{a_2}B$ | $A_bB$ | $AB_{a_1}$ | $AB_{a_2}$ | $AB_b$ | $A\|B$ |
|---|---|---|---|---|---|---|---|---|---|---|---|---|---|---|---|---|---|---|---|---|---|
| $AAA$ | $\cdot$ | $\varpi_{BA}$ | $\varpi_{BA}$ | | $\varpi_{BA}$ | | | | $c_A$ | $c_A$ | $c_A$ | | | | | | | | | | |
| $AAB$ | $\varpi_{AB}$ | $\cdot$ | | $\varpi_{BA}$ | | $\varpi_{BA}$ | | | | | | | | | | | | | | $c_A$ | |
| $ABA$ | $\varpi_{AB}$ | | $\cdot$ | $\varpi_{BA}$ | | | $\varpi_{BA}$ | | | | | | | | | | | | $c_A$ | | |
| $ABB$ | | $\varpi_{AB}$ | $\varpi_{AB}$ | $\cdot$ | | | | $\varpi_{BA}$ | | | | | | | $c_B$ | | | | | | |
| $BAA$ | $\varpi_{AB}$ | | | | $\cdot$ | $\varpi_{BA}$ | $\varpi_{BA}$ | | | | | | | | | | | $c_A$ | | | |
| $BAB$ | | $\varpi_{AB}$ | | | $\varpi_{AB}$ | $\cdot$ | | $\varpi_{BA}$ | | | | | | | | $c_B$ | | | | | |
| $BBA$ | | | $\varpi_{AB}$ | | $\varpi_{AB}$ | | $\cdot$ | $\varpi_{BA}$ | | | | | | | | | $c_B$ | | | | |
| $BBB$ | | | | $\varpi_{AB}$ | | $\varpi_{AB}$ | $\varpi_{AB}$ | $\cdot$ | | | | $c_B$ | $c_B$ | $c_B$ | | | | | | | |
| $A_{a_1}A$ | | | | | | | | | $\cdot$ | | | | | | $\varpi_{BA}$ | | | $\varpi_{BA}$ | | | $c_A$ |
| $A_{a_2}A$ | | | | | | | | | | $\cdot$ | | | | | | $\varpi_{BA}$ | | | $\varpi_{BA}$ | | $c_A$ |
| $A_bA$ | | | | | | | | | | | $\cdot$ | | | | | | $\varpi_{BA}$ | | | $\varpi_{BA}$ | $c_A$ |
| $B_{a_1}B$ | | | | | | | | | | | | $\cdot$ | | | $\varpi_{AB}$ | | | $\varpi_{AB}$ | | | $c_B$ |
| $B_{a_2}B$ | | | | | | | | | | | | | $\cdot$ | | | $\varpi_{AB}$ | | | $\varpi_{AB}$ | | $c_B$ |
| $B_bB$ | | | | | | | | | | | | | | $\cdot$ | | | $\varpi_{AB}$ | | | $\varpi_{AB}$ | $c_B$ |
| $A_{a_1}B$ | | | | | | | | | $\varpi_{AB}$ | | | $\varpi_{BA}$ | | | $\cdot$ | | | | | | |
| $A_{a_2}B$ | | | | | | | | | | $\varpi_{AB}$ | | | $\varpi_{BA}$ | | | $\cdot$ | | | | | |
| $A_bB$ | | | | | | | | | | | $\varpi_{AB}$ | | | $\varpi_{BA}$ | | | $\cdot$ | | | | |
| $AB_{a_1}$ | | | | | | | | | $\varpi_{AB}$ | | | $\varpi_{BA}$ | | | | | | $\cdot$ | | | |
| $AB_{a_2}$ | | | | | | | | | | $\varpi_{AB}$ | | | $\varpi_{BA}$ | | | | | | $\cdot$ | | |
| $AB_b$ | | | | | | | | | | | $\varpi_{AB}$ | | | $\varpi_{BA}$ | | | | | | $\cdot$ | |
| $A\|B$ | | | | | | | | | | | | | | | | | | | | | $\cdot$ |

Note.— $\varpi_{AB} = 4M_{AB}/\theta_B = m_{AB}/\mu$ and $\varpi_{BA} = 4M_{BA}/\theta_A = m_{BA}/\mu$ are mutation-scaled migration rates, and $c_A = 2/\theta_A$ and $c_B = 2/\theta_B$ are the coalescent rates, when one time unit is the expected time to accumulate one mutation per site. The state of the chain is given by the population IDs ($A$ or $B$) and sequence IDs. For example the initial state $A_{a_1}A_{a_2}B_b$ means that the three sequences $a_1, a_2$, and $b$ are from populations $A$, $A$, and $B$, respectively. States with three sequences are abbreviated, with the three sequences assumed to be in the order $a_1, a_2, b$ so that the sequence IDs are suppressed. Thus $A_{a_1}A_{a_2}B_b$ is '$AAB$'. State $A_{a_1a_2}B_b$ means that two sequences remain in the sample, with the ancestor of sequences $a_1$ and $a_2$ is in population $A$ while sequence $b$ is in population $B$. This is abbreviated '$AB_b$', with the sequence ID '$a_1a_2$' suppressed. '$A|B$' is an absorbing state in which only one sequence remains in the sample, in either $A$ or $B$, after two coalescent events have occurred. From Leaché *et al.* (2019).
